# Comparative Huntington and Parkinson Disease mRNA Analysis Reveals Common Inflammatory Processes

**DOI:** 10.1101/139451

**Authors:** Adam Labadorf, Seung Hoan Choi, Richard H Myers

## Abstract

Huntington’s and Parkinson’s Diseases (HD and PD) are neurodegenerative disorders that share some pathological features but are disparate in others. For example, while both diseases are marked by aberrant protein aggregation in the brain, the specific proteins that aggregate and types of neurons affected differ. A better understanding of the molecular similarities and differences between these two diseases may lead to a more complete mechanistic picture of both the individual diseases and the neurodegenerative process in general. We sought to characterize the common transcriptional signature of HD and PD as well as genes uniquely implicated in each of these diseases using mRNA-Seq data from post mortem human brains in comparison to neuropathologically normal controls. The enriched biological pathways implicated by HD differentially expressed genes show remarkable consistency with those for PD differentially expressed genes and implicate the common biological processes of neuroinflammation, apoptosis, transcriptional dysregulation, and neuron-associated functions. Comparison of the differentially expressed (DE) genes highlights a set of consistently altered genes that span both diseases. In particular, processes involving nuclear factor kappa-light-chain-enhancer of activated B cells (NFkB) and transcription factor cAMP response element-binding protein (CREB) are the most prominent among the genes common to HD and PD. When the combined HD and PD data are compared to controls, relatively few additional biological processes emerge as significantly enriched suggesting that most pathways are independently seen within each disorder. Despite showing comparable numbers of DE genes, DE genes unique to HD are enriched in far more coherent biological processes than the DE genes unique to PD, suggesting that PD may represent a more heterogeneous disorder.

## Introduction

Transcriptional dysregulation has been observed in both Huntington’s disease (HD) and Parkinson’s disease (PD)^1,2^. Transcription, neuroinflammation, and developmental processes have been shown to be dysregulated in the brains of HD individuals^3^, while inflammation and mitochondrial dysfunction were observed to be altered in the brains of PD individuals^4^. However, a systematic comparison of the transcriptional signatures of HD and PD has not been performed to date, and those genes and biological processes common to both diseases, if any, remain to be determined. To address this question, we sought to identify genes that are consistently differentially expressed (DE) in the post-mortem brains of HD and PD human subjects compared to neuropathologically normal control brains using mRNA-Seq. We hypothesize that common altered genes and pathways in HD and PD will elucidate the mechanistic underpinnings of the neurodegenerative process.

This study presents the results of a comparison of DE genes for each of HD and PD versus controls (C) analyzed separately. In addition, in order to identify consistent effects with lower effect size across diseases, an analysis was performed where the HD and PD datasets are concatenated as a single category, neurodegenerative disease (ND), and compared with C. DE genes are determined using a tailored form of logistic regression as described in^5^, which improves control of type I errors when compared to negative binomial based DE detection methods.

## Materials and Methods

### Sample collection, processing, and sequencing

The HD, PD, and C samples used in this study are those previously described in our past work^3,4^. Nine additional HD brain samples were included in this study beyond those in^3^, including two HD gene positive asymptomatic individuals, obtained from the Harvard Brain Tissue Resource Center. All samples underwent the same tissue dissection and RNA extraction sample preparation protocol performed by the same individuals. Briefly, RNA was extracted from the prefrontal cortex of post-mortem brains of HD and PD subjects, as well as neuropathologically normal controls. RNA was poly-A selected and subjected to mRNA sequencing on the Illumina HiSeq 2000 platform. Sample statistics are contained in Table 1. See^3^ and^4^ for more detailed information about sample preparation.

**Table 1.** Sample statistics. SD is standard deviation. PMI is Post Mortem Interval. RIN is RNA Integrity Number of the RNA extracted from each sample as determined by the Agilent Bioanalyzer instrument.

| Sample type | N | Mean (SD) Age at Death | Mean (SD) PMI | Mean (SD) RIN |
| --- | --- | --- | --- | --- |
| HD | 29 | 60.5 (11.4) | 16.4 (7.8) | 7.1 (1.2) |
| PD | 29 | 77.5 (8.9) | 11.1 (9.7) | 7.0 (0.7) |
| Control | 49 | 68.6 (15.8) | 14.6 (9.5) | 7.8 (0.7) |

### mRNA-Seq data processing

Each FASTQ file containing mRNA sequences was first trimmed for sequence quality using the sickle software package^6^ with default arguments. The resulting short reads were aligned against the hg38 build of the human reference genome using the STAR aligner v2.4.0h1^7^ with permissive multimapping parameters (200 maximum alignments —outFilterMultimapNmax 200) and otherwise parameter values suggested in the STAR manual. Multimapped reads were assigned unique alignment locations using the ORMAN software package^8^. Aligned reads were counted against GENCODE v21 annotation^9^ using the HTSeq package v0.6.1p1^10^. Read counts for all samples were normalized using the method described in the DESeq2 package v1.10.1^11^ and outlier counts were trimmed using the strategy described in^3^. Since the original mRNAs were poly-A selected, only genes with biotypes known to be polyadenylated `(i.e. protein_coding, lincRNA, processed_transcript, sense_intronic, sense_overlaping, IG_V_gene, IG_D_gene, IG_J_gene, IG_C_gene, TR_V_gene, TR_D_gene, TR_J_gene, and TR_C_gene)` as annotated by Ensembl BioMart^12^ downloaded on May 27th, 2015. To avoid spurious results due to low abundance, genes were further filtered if more than half of the ND or C samples had zero counts.

### Differential expression and assessment of batch effects

DE genes were identified using Firth’s logistic (FL) regression^13,14^ applied to mRNA-Seq data as described in^5^. Briefly, in contrast to negative binomial regression models like edgeR^15^ and DESeq2^11^, this method models a binomial status variable (e.g. case vs control) as a function of gene counts and any other potentially confounding variables (RIN value, PMI, etc.). Classical logistic regression has historically not been used to determine DE genes because of the so-called “complete separation” problem, where model parameter estimation fails when there is perfect or nearly perfect separation of a predictor with respect to a binomial variable (e.g. one condition has extremely high read counts and the other has very low read counts). FL regression addresses this issue by using a modified likelihood function to enable reliable parameter estimation, and has other statistical advantages with respect to type I error rates and power. Note the DE statistic from FL regression is log odds ratio (LOR) of case versus controls, that is, positive LOR indicates greater mRNA abundance in case and negative LOR indicates greater abundance in control. All reported p-values are Benjamini-Hochberg (BH) adjusted^16^ unless noted otherwise. See^5^ for further information on this method applied to mRNA-Seq data.

The data sets in this study were sequenced in five separate batches. To evaluate whether there was evidence of systemic batch effects confounding the identification of DE genes, we ran separate Firth DE models with and without a categorical variable representing batches and compared the beta estimates using Spearman correlation. Beta estimate ranks were highly correlated for HD vs C (ρ = 0.84, *p* ⋘ 0.001), PD vs C (ρ = 0.99, p ⋘ 0.001), and ND vs C (ρ = 0.97, *p* ⋘ 0.001). We therefore did not include a batch variable in the DE models.

### Identification of ND DE genes and enriched genesets

DE genes were identified as those with BH adjusted p-values < 0.01 from the Firth’s logistic regression models of HD vs C, PD vs C, and ND vs C models, yielding three independent DE gene lists. Read counts for each gene were scaled to have a mean of zero and standard deviation of one to obtain standardized regression coefficients, which makes coefficients comparable across genes. All controls were used in each model. Gene set enrichment analysis was performed on each gene list ranked by read counts beta coefficient using the GSEA^17^ implementation in the DOSE software package^18^ against the MsigDB Canonical Pathway (C2) geneset database^17^. GSEA enrichment was computed using the complete list of genes irrespective of significance ranked by standardized beta coefficient of the count variable. The robust rank aggregation (RRA) algorithm^19^ was used to identify individual genes that were consistently altered across these gene lists. Briefly, RRA is a probabilistic, non-parametric, rank-based method for detecting genes ranked consistently better than expected under the null hypothesis of uncorrelated inputs in the presence of noise and outliers. The genes identified as most significant by RRA are the most likely to be implicated in the general ND phenotype.

In addition to producing independent HD and PD DE gene lists, we sought to functionally characterize the genes that are uniquely significant to each disease as well as those in common. To accomplish this, the DE genes from HD and PD were intersected, partitioning the genes into HD-specific, PD-specific, and DE genes common to the two gene lists. Each of these partitioned gene lists were then subjected to gene set enrichment on the MsigDB Canonical Pathway (C2), Transcription Factor Targets (C3), and Gene Ontology (C5) gene set databases^17^ using a hypergeometric test.

## Results

Firth’s logistic (FL) regression identified 2427, 1949, and 4843 significantly DE genes for HD, PD, and ND, respectively, at q-value < 0.01. Gene set enrichment analysis of MsigDB C2 gene sets identified 226, 199, and 250 gene sets significantly enriched at q-value < 0.05 for HD, PD, and ND, respectively. Due to the large number of DE genes in each dataset, we focus exclusively on the GSEA enrichment results here. Complete DE gene list and gene set enrichment statistics for HD, PD, and ND are included in Supplemental File 1.

There was a high degree of overlap between the significantly enriched gene sets of HD and PD. 145 gene sets were significantly enriched in both DE gene lists, while 81 and 54 gene sets were uniquely significant in HD and PD, respectively. When a pathway was enriched in more than one list, the pathway was always, without exception, enriched in the same direction, either positively (genes are more abundant in disease) or negatively (genes are less abundant in disease). There were 24 gene sets uniquely significant in ND. Figure 1 depicts the enriched gene sets grouped by high-level biological category for HD, PD, and ND.

**Figure 1.**
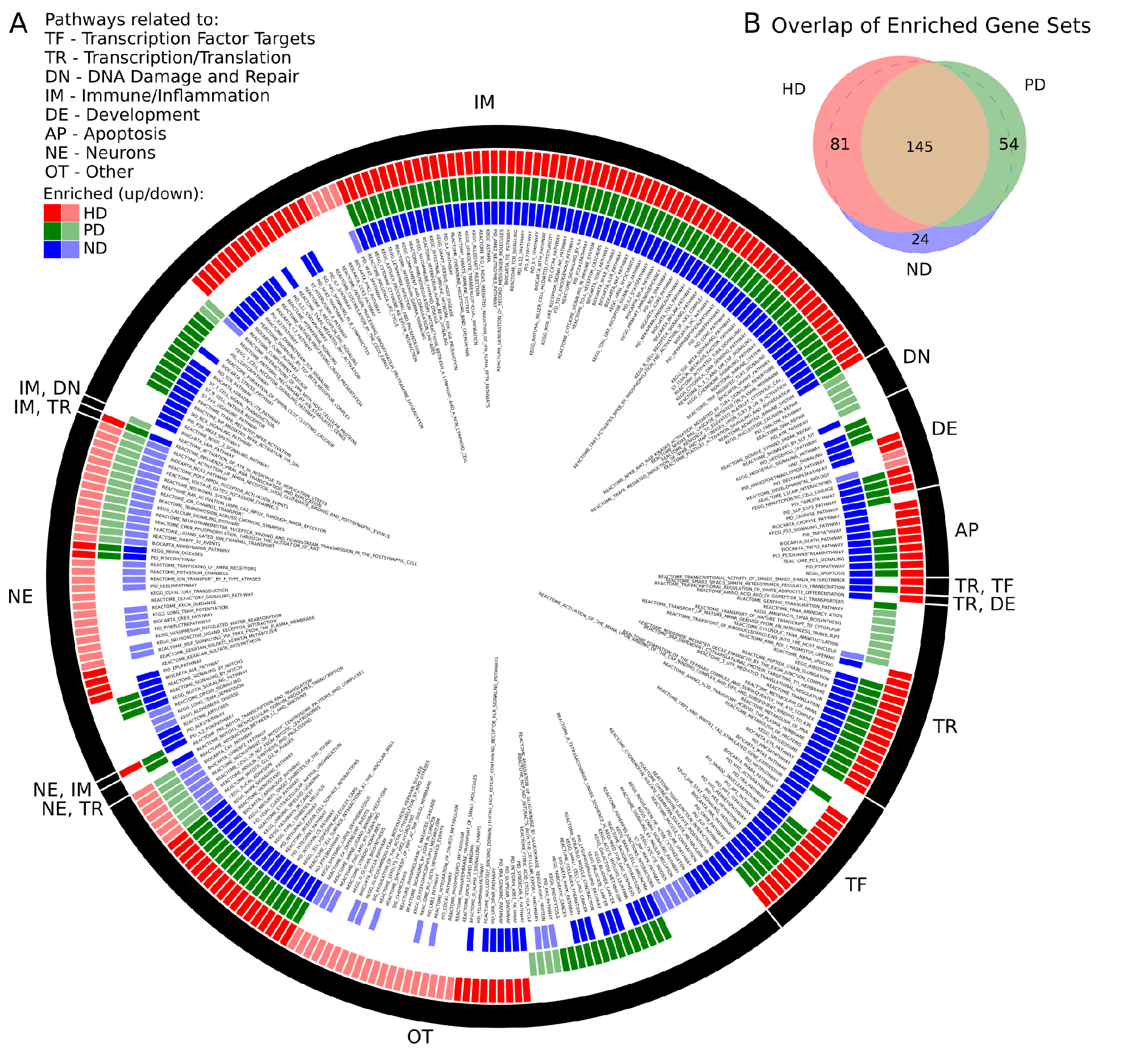
A) Significantly enriched MsigDB C2 Canonical Pathway gene sets for HD, PD, and ND identified by GSEA. Each colored ring segment corresponds to a single enriched gene set. Red (outer), green (middle), and blue (inner) segmented rings indicate whether the HD, PD, or ND DE gene lists, respectively, were significantly enriched for the gene set. Dark and light colored segments indicate up and down regulation (positive, negative GSEA normalized enrichment score), respectively. Black ring around exterior groups gene sets into high level categories as indicated by the two letter code. Gene set name is listed in interior of rings. A tabular form of the data underlying the figure is in Supplemental File 1. B) Venn diagram illustrating overlap of significantly enriched gene sets for HD, PD, and ND. All but 24 of the ND significant gene sets were significantly enriched in either HD, PD, or both.

We make several observations of Figure 1 A. First, the plurality of enriched gene sets across all three data sets are related to immune processes (IM) and are with few exceptions positively enriched. Pathways related to neuronal processes (NE) are largely negatively enriched and there is a subset of these gene sets that are exclusively enriched in HD. With the exception of DNA damage, all remaining biological categories are represented for both HD and PD. DNA damage related pathways (DN) are unique to the PD dataset and are negatively enriched. Multiple apoptosis (AP), developmental (DE), transcription/translation (TR), and transcription factor target (TF) gene sets are also enriched in all three gene lists.

There were 83 gene sets that did not fit cleanly into a single category (OT), which notably include pathways related to endocytosis, signaling, cellular adhesion and extracellular matrix, glycans, and metabolism. A tabular form of the data underlying Figure 1 A is in Supplemental File 1.

RRA identified 1353 genes with a score < 0.01. The top ten genes identified by RRA as most highly ranked across all three gene lists are reported in Table 2. The rank of each gene in the individual gene lists are also reported in the table, showing that most genes are relatively highly ranked across all three studies as expected. The list of all significant genes identified by RRA analysis is included in Supplemental File 1.

**Table 2.** Top ranked RRA genes. RRA Score can be thought of as a p-value. The remaining columns contain the rank of the corresponding gene in each individual gene list.

| Symbol | Description | RRA Rank | RRA Score | HD Rank | PD Rank | ND Rank |
| --- | --- | --- | --- | --- | --- | --- |
| ENSG00000272403 | no description | 1 | $4.39 \times 10^{-9}$ | 25 | 15 | 1 |
| SPR | sepiapterin reductase | 2 | $1.13 \times 10^{-7}$ | 70 | 38 | 5 |
| DDIT4 | DNA-damage-inducible transcript 4 | 3 | $1.27 \times 10^{-7}$ | 74 | 60 | 22 |
| TRIP10 | thyroid hormone receptor interactor 10 | 4 | $1.87 \times 10^{-7}$ | 84 | 59 | 7 |
| TNFRSF10D | tumor necrosis factor receptor superfamily, member 10d | 5 | $2.30 \times 10^{-7}$ | 90 | 55 | 20 |
| PRMT6 | Protein arginine methyltransferase 6 | 6 | $2.54 \times 10^{-7}$ | 29 | 93 | 10 |
| GPSM3 | G-protein signaling modulator 3 | 7 | $2.62 \times 10^{-7}$ | 81 | 98 | 11 |
| GPCPD1 | Glycerophosphocholine phosphodiesterase 1 | 8 | $2.79 \times 10^{-7}$ | 13 | 96 | 2 |
| GPR4 | G-protein-coupled receptor 4 | 9 | $2.97 \times 10^{-7}$ | 98 | 75 | 24 |
| NFKBIA | Nuclear factor of kappa light polypeptide gene enhancer in B-cells inhibitor, Alpha | 10 | $3.35 \times 10^{-7}$ | 11 | 103 | 3 |

The most consistently ranked gene is RP1-93H18.7 (ENSG00000272403.1), a lncRNA, which was removed from Ensembl starting at version GRCh38.p2, but shows consistent transcription in these data. This gene is directly downstream of the gene DSE (dermatan sulfate epimerase) which is also DE in both HD and PD, is involved in embryonic development^20,21^, and has been related to the immune response in cancer patients^22^. Deficiencies in the second ranked gene, SPR (sepiapterin reductase), have been linked to DOPA-responsive dystonia^23^ and previously implicated in PD^24^. The third gene, DDIT4 (DNA-Damage-Inducible Transcript 4), is a multi-functional gene which, via its inhibition of the mammalian target of rapamycin complex 1 (mTORC1), regulates in cell growth, proliferation, and survival^25^, controls p53/TP53-mediated apoptosis in response to DNA damage^26,27^, and plays a role in neurodegeneration, neuronal death and differentiation, and neuron migration during embryonic brain development^28–31^. TRIP10 (thyroid hormone receptor interactor 10), another multi-functional gene, is involved in insulin signaling^32^, endocytosis^33^, and structures specific to monocyte-derived cells^34^. TNFRSF10D (Tumor Necrosis Factor Receptor Superfamily, Member 10d, Decoy With Truncated Death Domain) inhibits certain types of apoptosis and may play a role in NfkB pathway^35^.

Figure 2 illustrates the differences in normalized counts for the top genes identified by RRA. With the exception of (12) PITX1, which is driven entirely by HD, all top genes demonstrate substantial differences between both disease conditions and control.

**Figure 2.**
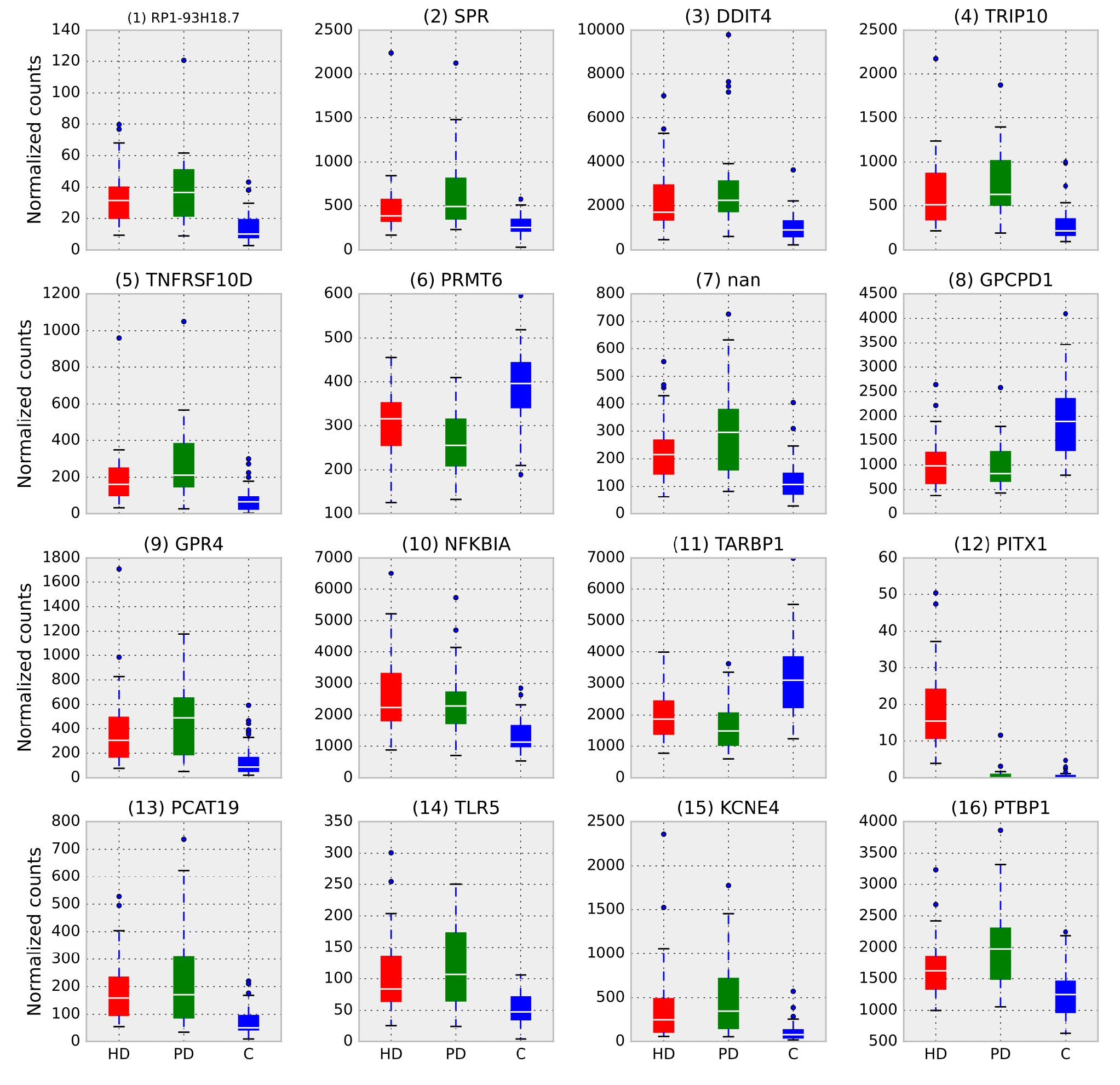
Box plots of normalized counts for top RRA genes split by condition, RRA rank is in parenthesis. Whiskers extend to 25th and 75th percentile counts, white bars are median counts. With the exception of (12) PITX1, which is driven entirely by HD, all top genes demonstrate substantial differences between both disease conditions and control.

Finally, we examined the significant DE genes from HD and PD for intersection. Figure 3 illustrates the overlap of DE genes between diseases and describes gene set enrichment results for the intersection. Figure 4 contains the enrichment results for the HD unique genes.

**Figure 3.**
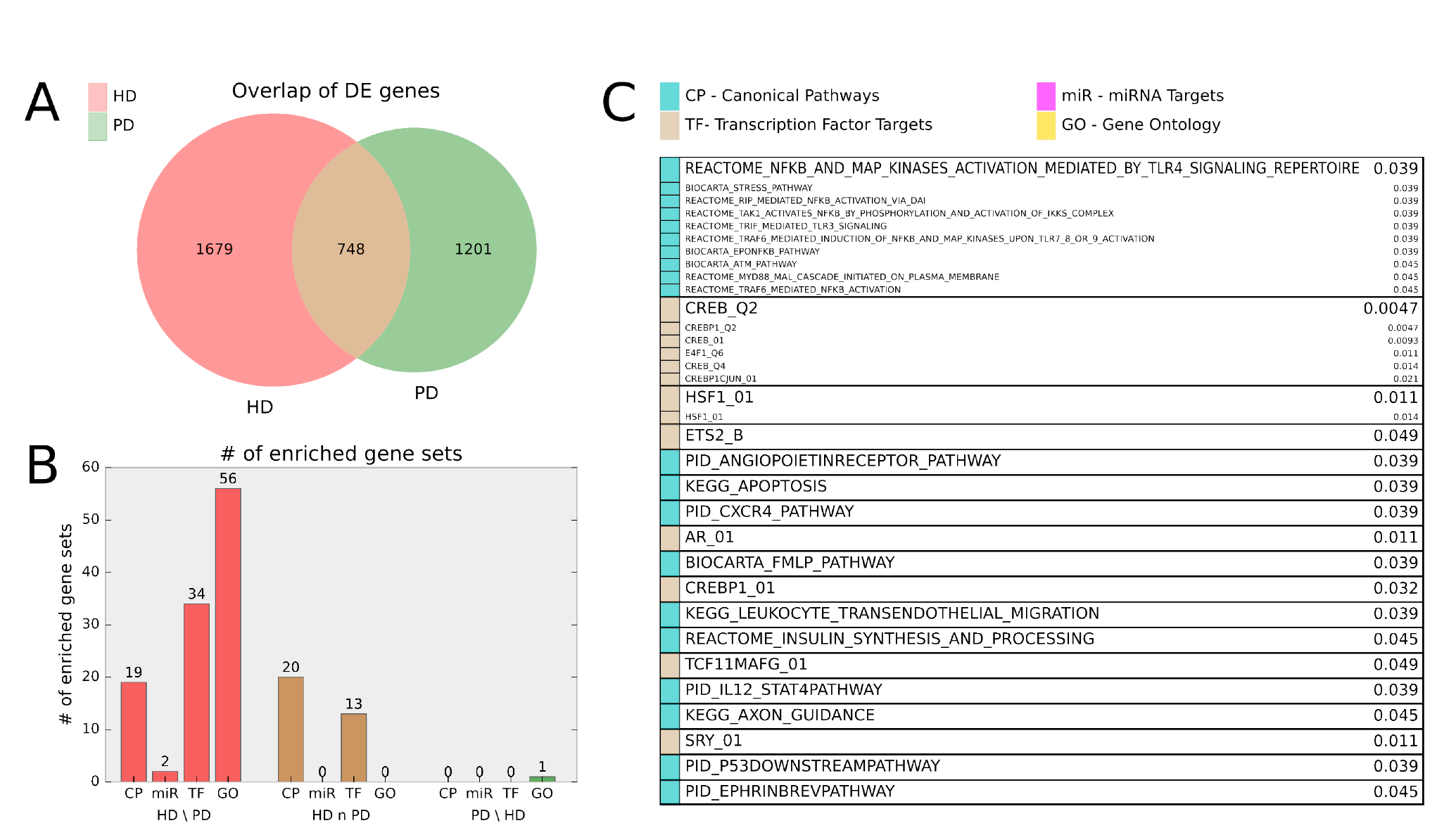
A) Venn diagram of HD and PD DE gene list intersection for DE genes adjusted pi0.01. B) Bar chart indicating number of MsigDB C2 Canonical Pathway (CP), C3 miRNA Targets (miR), C3 Transcription Factor Targets (TF), and C5 Gene Ontology (GO) gene sets enriched for the HD unique (HD \PD), intersection (HD n PD), and PD unique (PD \HD) genes. For HD \PD enrichment, 17 redundant or uninformative GO gene sets and 7 TF gene sets for motifs with unknown transcription factors were omitted from the figure results but are included in Supplemental File 1 Table 1. C) Gene sets enriched for the intersection genes (HD n PD). Adjusted p-values are listed beside gene set name and the originating gene set (CP, miR, TF, or GO) are indicated by color. Gene sets that are grouped into boxes share more than 20% of their DE genes and are therefore listed together.

**Figure 4.**
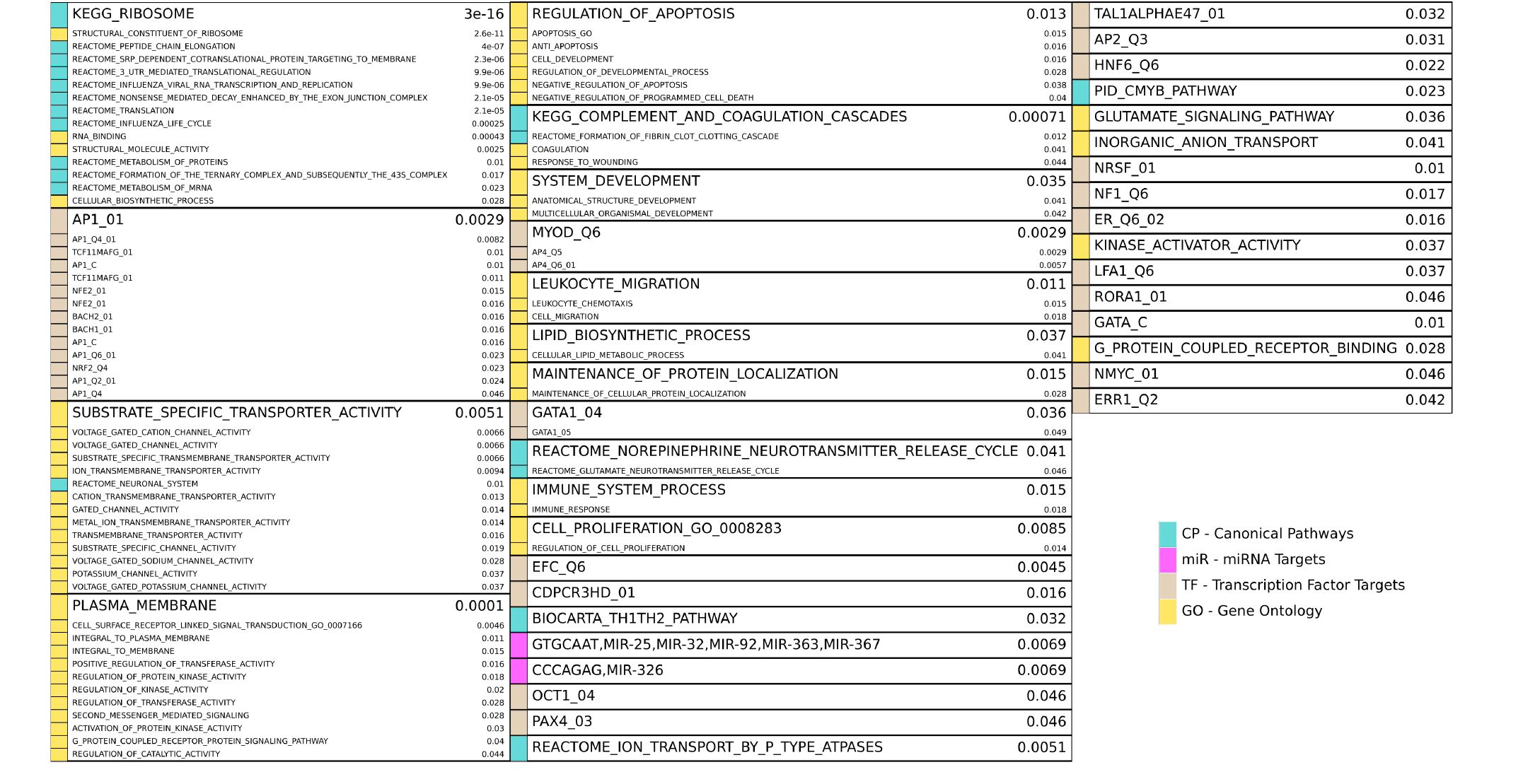
Enriched gene sets for the HD unique (HD PD) genes from Figure 3 A and reported similarly as in Figure 3 C. Note 17 redundant or uninformative GO gene sets and 7 TF gene sets for motifs with unknown transcription factors were omitted from the figure results but are included in Supplemental File 1 Table 1.

HD specific enrichments are shown in Figure 4, where a broad spectrum of biological processes are implicated in the HD-unique DE genes. The most striking enriched gene set is KEGG_RIBOSOME, with many other related gene sets involved in translation and molecular metabolism similarly enriched. Multiple gene sets that share > 20% of their DE genes are associated with Jun Proto-Oncogene (AP1), BTB And CNC Homology 1, Basic Leucine Zipper Transcription Factor 1 and 2 (BACH1, BACH2), and NRF2/TCF11 are also implicated. Other strongly implicated biological processes are ion channel activity, plasma membrane and signaling, apoptosis, immune system and inflammatory processes, developmental genes, neuron-related signaling pathways, many transcription factors, and two families of miRNAs.

## Discussion

RNA-sequencing was performed in 29 HD, 29 PD and 49 control prefrontal cortex (BA9) samples to assess the common and unique transcriptional profiles of these two protein aggregation related neurodegenerative diseases. Firth’s logistic (FL) regression identified 2427, 1949, and 4843 significantly DE genes for HD, PD, and ND, respectively, at q-value < 0.01. Gene set enrichment analysis of MsigDB C2 gene sets identified 226, 199, and 250 gene sets significantly enriched at q-value < 0.05 for HD, PD, and ND, respectively. Gene sets related to immune processes and inflammatory pathways, were highly enriched for both diseases. Notably, Figure 1 shows that the overwhelming majority of enriched biological pathways are common to both diseases and that they are invariably perturbed in the same direction in both diseases. To the authors’ knowledge, this study presents the first comprehensive comparative analysis of DE gene expression from HD, PD, and ND in post-mortem human brains assessed with mRNA-Seq. In one previous study, Capurro et al analyzed HD and PD microarray data using a cell-type deconvolution method to identify cell-type specific differences in gene expression between cases and controls for both diseases^36^. Only one gene from the study, doublecortin-like kinase 1 (DCLK1), was found to be differentially expressed in both HD and PD cortex, and this gene also appears as DE in common between diseases in the analysis presented here.

The comparison of HD and PD in particular is motivated by the fact that these diseases can be viewed as mirror-images of each other. GABAergic medium spiny interneurons, which compose most of the neurons in the striatum and selectively die in HD but are spared in PD, project directly into the substantia nigra and coordinate motor activity throughout the brain via dopamine-induced signaling^37^. Dopaminergic neurons in the substantia nigra, on the other hand, which also are important in coordinating motor activity as well as arousal, reinforcement, and reward^38^, selectively degenerate in PD but are spared in HD. It was observed in a study of 523 HD subjects that the incidence of PD in this cohort was lower than that of the general population, though both HD and PD individuals develop Alzheimer’s disease at the same rate^39^, suggesting the selective death of medium spiny neurons might be neuroprotective of dopaminergic neuron death. Given the intimate neurological link between the affected neurons in HD and PD, and the mutual exclusivity of their degeneration, this comparison poses a very interesting contrast to identify common responses to neurodegeneration that are not confounded by neuron type. Unfortunately, a direct comparison of neurons in these regions of post-mortem human brains is not possible, precisely due to this mirror-image pathology. The choice of the BA9 brain region is motivated by the fact that, due to degeneration, the affected neurons are largely missing from the striatum and substantia nigra in HD and PD, respectively, whereas BA9 is generally free of involvement in both diseases^39–41^. Because the primarily affected neurons in HD and PD do not exist in BA9, the biological processes implicated by this analysis may likely represent the response to, rather than direct cause of, the respective diseases. Nonetheless, the remarkable consistency between HD and PD observed in this analysis points to important mechanisms that further our understanding of neurodegenerative disease as a general process.

The biological processes implicated by DE gene lists identified from each condition separately are compellingly similar. From Figure 1, we see that the majority of enriched biological pathways are common and that they are invariably perturbed in the same direction in both diseases. Furthermore, combining HD and PD data into an ND condition does not yield significantly more novel biological insights. This remarkable consistency between the pathway enrichment results suggest that the underlying molecular responses to neurodegeneration in HD and PD may be more similar than they are different, despite their different pathological underpinnings. Of particular significance is the strong positive enrichment of immune and inflammatory pathways, which have been convincingly implicated in both diseases separately^3,42–45,45–48^, but the compelling similarity of these signatures between HD and PD revealed by this analysis has not been illustrated to date.

The negative regulation of neuron-related pathways is also noteworthy, since the BA9 brain region, from which these samples are derived, is not known to be heavily involved in either of these diseases. Despite the lack of clear and consistent neurodegeneration in this brain region, the widespread biological pathways shown to be affected in this analysis strongly suggest neurons in BA9 do indeed experience a common set of effects in the neuropathology for HD and PD.

Many of the individual genes identified by RRA as most consistently different in HD, PD, and ND have previously been the focus of studies in neurodegeneration. The second highest ranked gene SPR has been the focus of significant study in PD and is related to the PARK3 gene locus^24,49^, but has not been previously implicated in HD. Inhibition of DNA-damage inducible transcript 4 (DDIT4/RTP801/REDD1) has been associated with neuroprotection in PD models and patients^31^ and is involved with mutant Huntingtin-induced cell death^50^. Thyroid hormone receptor interactor 10 (TRIP10) has been shown to interact directly with mutant huntingtin^51^, and while it is not known to play a role in PD pathology, its elevated mRNA abundance in these PD samples suggest it may indeed be implicated. Other top genes have also been implicated in neurodegeneration: tumor necrosis factor receptor superfamily 10D (TNFRSF10D)^52,53^, protein arginine methyltransferase 6 (PRMT6)^54^, and toll-like receptor 5 (TLR5)^55^. Further investigation of this list of genes is likely to yield novel insights into the mechanisms of neurodegeneration.

The intersection of DE genes between HD and PD also affords important insight into genes relevant to fundamental neurodegenerative processes. Most notably, pathways related to NFkB and transcriptional targets of CREB are prominent in the enrichment results. The NFkB pathway is prominent in both HD^56,57^ and PD pathology^58,59^ through its central role in inflammatory signaling. CREB is directly targeted by brain derived-neurotrophic factor (BDNF)^60^, an important trophic factor in the brain. Both BDNF^61^, and CREB^5,62^ have been directly implicated in HD pathology, while CREB is also believed to play a critical role in dopamine receptor-mediated nuclear signaling^63^, and disruption of the protein’s function leads to neurodegeneration^64,65^. The specific inflammation-related gene sets (HSF1 transcription factor targets, CXCR4, IL12) suggests there is some specificity in the aspects of the pan-neurodegenerative neuroimmune response. Recent studies in both HD and PD have focused on the role of insulin sensitivity and metabolism in patients^66–68^, supporting the role of insulin synthesis as an enriched biological pathway in the common gene list. While the enrichment of apoptosis-related pathways was not surprising, pathways related to angiopoietin, ephrin, and axon guidance suggest that biological processes related to neuronal plasticity are active in both of these diseases and may even indicate that neuroprotective or neuroregenerative processes are a component of the neurodegenerative response.

These data also point to compelling differences between HD and PD. Interestingly, two groups of genes, DNA damage and repair and tRNA related processes, seem to be uniquely perturbed and negatively enriched in PD. The DNA damage and repair gene set enrichment may be a reflection of mitochondrial DNA damage. In PD, dopaminergic neurons of the substantia nigra (though not cortical neurons) were found to be particularly vulnerable to mitochondrial DNA damage^69^, and Lewy body pathology, the histological hallmark of PD, is associated with mitochondrial DNA damage^70^. More generally, mitochondrial DNA damage and oxidative stress are associated with several neurodegenerative diseases including PD, Alzheimer’s disease^71^, and ALS^72^. There is evidence supporting the involvement of aminoacyl tRNA synthetases in neurological disease, including ALS, leukoencephalopathy, and PD^73^.

In HD, a number of uniquely perturbed gene sets related to glycan biosynthesis and metabolism are negatively regulated, and these pathways have not been previously implicated in HD. The 1687 HD-unique DE genes are enriched for many gene sets across a broad spectrum of biological processes, including mRNA and protein metabolism, ion channel activity, signaling and kinase activity, apoptosis, immune response, and development. Other, more specific gene sets related to a large number of transcription factors further support the observation of transcriptional dysregulation in HD^73,74^. The specificity of these enriched TF gene sets is quite striking, as the targeted DE genes appear to be largely disjoint between them, suggesting potential, specific causes of the dysregulated transcriptional effects in HD. The enrichment of two miRNA families are also particularly relevant in light of recent reports of miRNA dysregulation in HD^75,76^.

It is interesting to note the disparity in enrichment between the HD and PD unique DE genes. Though the numbers of unique DE genes are comparable, the large number of enriched gene sets in HD stands in sharp contrast to the almost total absence of enrichment in PD. This result implies that the DE genes in HD are more consistently related to one another than in PD. One possible, and potentially important explanation for this is that HD is a much more homogeneous disease than PD. It is well established that PD has a significant sporadic component^77^, caused by a combination of genetic and environmental factors. The relative heterogeneity of PD may make finding consistently effective treatments difficult, and the absence of biological enrichment in specific pathways, other than those common to both diseases, from this analysis may be a reflection of an underlying molecular basis for this effect. It may be that, given sufficient sample size, coherent subgroups of patients may be identified by examining patterns in their gene expression using datasets such as those analyzed here. On the other hand, despite extensive molecular characterization of HD, effective, widely available therapies for HD have remained elusive despite the relative homogeneity of the disease process among HD patients.

These findings have important implications on our understanding of the neurodegenerative disease process. The significant involvement of the inflammatory pathways in both diseases in an area not thought to be directly involved in disease pathogenesis suggests the response to neurodegeneration is widespread throughout the brain. NFkB in particular appears to be a major player, which is well supported in the HD and PD literature. It is unclear whether the neuroinflammatory response is protective, deleterious, or both from these data, but investigation into the role these processes play, and the potential therapeutic value of modulating them, should be made a high priority.

## Author contributions statement

A.L. designed the implementation of the experiments, wrote all analysis code, and prepared the manuscript. S.C. provided guidance and critical feedback on the use of Firth logistic regression. R.M. conceived of the overall study, made available and processed the samples, and provided critical study and manuscript feedback.

## Additional information

Raw FASTQ reads for these samples are available under GEO accession numbers GSE64810 and GSE68719. The code used to perform the statistical analysis and generate figures is available at https://bitbucket.org/adamlabadorf/hdvpd_mrnaseq.

## Acknowledgments

Supported by grants from US National Institutes of Health (R01-S076843), Characterization of the Role of Cyclin G-associated Kinase in Parkinson Disease, (R01-NS073947), Epigenetic Markers in Huntington’s Disease Brain, (R01-NS088538) An IPSc based platform for functionally assessing genetic and environmental Risk in PD, (U24-NS072026) National Brain and Tissue Resource for Parkinson’s Disease and Related Disorders, and the Jerry McDonald Huntington Disease Research Fund.

We would like to acknowledge the National Brain and Tissue Resource for Parkinson’s Disease and Related Disorders at Banner Sun Health Research Institute (NS072026), Sun City, Arizona, the Harvard Brain Tissue Resource Center McLean Hospital, Belmont, Massachusetts, and the Human Brain and Spinal Fluid Resource Center VA, West Los Angeles Healthcare Center, California for providing the brain samples used in these studies.

## References

1. Cha, J.-H. J. Transcriptional signatures in huntington’s disease. Prog. Neurobiol. 83, 228–248 (2007).

2. Elstner, M. et al. Expression analysis of dopaminergic neurons in parkinson’s disease and aging links transcriptional dysregulation of energy metabolism to cell death. Acta Neuropathol. 122, 75–86 (2011).

3. Labadorf, A. et al. RNA sequence analysis of human huntington disease brain reveals an extensive increase in inflammatory and developmental gene expression. PLoS One 10, e0143563 (2015).

4. Dumitriu, A. et al. Gene expression profiles in parkinson disease prefrontal cortex implicate FOXO1 and genes under its transcriptional regulation. PLoS Genet. 8, e1002794 (2012).

5. Choi, S. H. et al. Evaluation of logistic regression models and effect of covariates for case-control study in RNA-Seq analysis. BMC Bioinforma. 18, 91 (2017).

6. Joshi NA, Fass NJ. Sickle: A sliding-window, adaptive, quality-based trimming tool for FastQ files (2011).

7. Dobin, A. et al. STAR: ultrafast universal RNA-seq aligner. Bioinforma. 29, 15–21 (2013).

8. Dao, P. et al. ORMAN: Optimal resolution of ambiguous RNA-Seq multimappings in the presence of novel isoforms. Bioinforma. 30, 644–651 (2014).

9. Harrow, J. et al. GENCODE: The reference human genome annotation for the ENCODE project. Genome Res. 22, 1760–1774 (2012).

10. Anders, S., Pyl, P. T. & Huber, W. HTSeq - a python framework to work with high-throughput sequencing data. bioRxiv 002824 (2014).

11. Love, M. I., Huber, W. & Anders, S. Moderated estimation of fold change and dispersion for RNA-Seq data with DESeq2. bioRxiv 002832 (2014).

12. Kinsella, R. J. et al. Ensembl BioMarts: a hub for data retrieval across taxonomic space. Database 2011, bar030 (2011).

13. Firth, D. Bias reduction of maximum likelihood estimates. Biom. 80, 27–38 (1993).

14. Heinze, G. & Schemper, M. A solution to the problem of separation in logistic regression. Stat. Med. 21, 2409–2419 (2002).

15. Robinson, M. D., McCarthy, D. J. & Smyth, G. K. edger: a bioconductor package for differential expression analysis of digital gene expression data. Bioinforma. 26, 139–140 (2010).

16. Benjamini, Y. & Hochberg, Y. Controlling the false discovery rate: a practical and powerful approach to multiple testing. J. R. Stat. Soc. Ser. B Stat. Methodol. 57, 289–300 (1995).

17. Subramanian, A. et al. Gene set enrichment analysis: A knowledge-based approach for interpreting genome-wide expression profiles. Proc. Natl. Acad. Sci. U. S. A. 102, 15545–15550 (2005).

18. Yu, G., Wang, L.-G., Yan, G.-R. & He, Q.-Y. DOSE: an R/Bioconductor package for disease ontology semantic and enrichment analysis. Bioinforma. 31, 608–609 (2015).

19. Kolde, R., Laur, S., Adler, P. & Vilo, J. Robust rank aggregation for gene list integration and meta-analysis. Bioinforma. 28, 573–580 (2012).

20. Stachtea, X. N. et al. Dermatan Sulfate-Free mice display embryological defects and are neonatal lethal despite normal lymphoid and Non-Lymphoid organogenesis. PLoS One 10, e0140279 (2015).

21. Habicher, J. et al. Chondroitin / dermatan sulfate modification enzymes in zebrafish development. PLoS One 10, e0121957 (2015).

22. Mizukoshi, E. et al. Expression of chondroitin-glucuronate c5-epimerase and cellular immune responses in patients with hepatocellular carcinoma. Liver Int. 32, 1516–1526 (2012).

23. Wijemanne, S. & Jankovic, J. Dopa-responsive dystonia-clinical and genetic heterogeneity. Nat. Rev. Neurol. 11, 414–424 (2015).

24. Tobin, J. E. et al. Sepiapterin reductase expression is increased in parkinson’s disease brain tissue. Brain Res. 1139, 42–47 (2007).

25. Dennis, M. D., McGhee, N. K., Jefferson, L. S. & Kimball, S. R. Regulated in DNA damage and development 1 (REDD1) promotes cell survival during serum deprivation by sustaining repression of signaling through the mechanistic target of rapamycin in complex 1 (mTORC1). Cell. Signal. 25, 2709–2716 (2013).

26. Cam, M. et al. p53/TAp63 and AKT regulate mammalian target of rapamycin complex 1 (mTORC1) signaling through two independent parallel pathways in the presence of DNA damage. J. Biol. Chem. 289,4083–4094 (2014).

27. Vadysirisack, D. D., Baenke, F., Ory, B., Lei, K. & Ellisen, L. W. Feedback control of p53 translation by REDD1 and mTORC1 limits the p53-dependent DNA damage response. Mol. Cell. Biol. 31,4356–4365 (2011).

28. Romaní-Aumedes, J. et al. Parkin loss of function contributes to RTP801 elevation and neurodegeneration in parkinson’s disease. Cell Death Dis. 5, e1364 (2014).

29. Canal, M., Romaní-Aumedes, J., Martín-Flores, N., Perez-Fernandez, V. & Malagelada, C. RTP801/REDD1: a stress coping regulator that turns into a troublemaker in neurodegenerative disorders. Front. Cell. Neurosci. 8, >313 (2014).

30. Ota, K. T. et al. REDD1 is essential for stress-induced synaptic loss and depressive behavior. Nat. Med. 20, 531–535 (2014).

31. Malagelada, C. et al. RTP801/REDD1 regulates the timing of cortical neurogenesis and neuron migration. J. Neurosci. 31, 3186–3196 (2011).

32. Chang, L., Chiang, S.-H. & Saltiel, A. R. TC10a is required for Insulin-Stimulated glucose uptake in adipocytes. Endocrinol. (2013).

33. Feng, Y. et al. The cdc42-interacting protein-4 (CIP4) gene knock-out mouse reveals delayed and decreased endocytosis. J. Biol. Chem. 285, 4348–4354 (2010).

34. Linder, S., Hüfner, K., Wintergerst, U. & Aepfelbacher, M. Microtubule-dependent formation of podosomal adhesion structures in primary human macrophages. J. Cell Sci. 113 Pt 23, 4165–4176 (2000).

35. Degli-Esposti, M. A. et al. The novel receptor TRAIL-R4 induces NF-kappaB and protects against TRAIL-mediated apoptosis, yet retains an incomplete death domain. Immun. 7, 813–820 (1997).

36. Capurro, A., Bodea, L.-G., Schaefer, P., Luthi-Carter, R. & Perreau, V. M. Computational deconvolution of genome wide expression data from parkinson’s and huntington’s disease brain tissues using population-specific expression analysis. Front. Neurosci. 8, 441 (2014).

37. Yager, L. M., Garcia, A. F., Wunsch, A. M. & Ferguson, S. M. The ins and outs of the striatum: role in drug addiction. Neurosci. 301, 529–541 (2015).

38. Schultz, W. Multiple dopamine functions at different time courses. Annu. Rev. Neurosci. 30, 259–288 (2007).

39. Hadzi, T. C. et al. Assessment of cortical and striatal involvement in 523 huntington disease brains. Neurol. 79, 1708–1715 (2012).

40. Braak, H. et al. Staging of brain pathology related to sporadic parkinson’s disease. Neurobiol. Aging 24, 197–211 (2003).

41. Halliday, G. M., Del Tredici, K. & Braak, H. Critical appraisal of brain pathology staging related to presymptomatic and symptomatic cases of sporadic parkinson’s disease. J. Neural Transm. Suppl. 99–103 (2006).

42. Kwan, W. et al. Mutant huntingtin impairs immune cell migration in huntington disease. J. Clin. Invest. 122, 4737–4747 (2012).

43. Crotti, A. et al. Mutant huntingtin promotes autonomous microglia activation via myeloid lineage-determining factors. Nat. Neurosci. 17, 513–521 (2014).

44. Ellrichmann, G., Reick, C., Saft, C. & Linker, R. A. The role of the immune system in huntington’s disease. J. Immunol. Res. 2013, e541259 (2013).

45. Dexter, D. T. & Jenner, P. Parkinson disease: from pathology to molecular disease mechanisms. Free. Radic. Biol. Medicine 62, 132–144 (2013).

46. Dobbs, R. J. et al. Association of circulating TNF-α and IL-6 with ageing and parkinsonism. Acta Neurol. Scand. 100, 34–41 (1999).

47. Jenner, P. Oxidative stress in parkinson’s disease. Ann. Neurol. 53 Suppl 3, S26–36; discussion S36-8 (2003).

48. Allen Reish, H. E. & Standaert, D. G. Role of α-Synuclein in inducing innate and adaptive immunity in parkinson disease. J. Park. Dis. 5, 1–19 (2015).

49. Sharma, M. et al. Role of sepiapterin reductase gene at the PARK3 locus in parkinson’s disease. Neurobiol. Aging 32, 2108.e1–5 (2011).

50. Martín-Flores, N. et al. RTP801 is involved in mutant Huntingtin-Induced cell death. Mol. Neurobiol. (2015).

51. Holbert, S. et al. Cdc42-interacting protein 4 binds to huntingtin: neuropathologic and biological evidence for a role in huntington’s disease. Proc. Natl. Acad. Sci. U. S. A. 100, 2712–2717 (2003).

52. Lopez-Gomez, C. et al. TRAIL/TRAIL receptor system and susceptibility to multiple sclerosis. PLoS One 6, e21766 (2011).

53. Frenkel, D. A new TRAIL in alzheimer’s disease therapy. Brain 138, 8–10 (2015).

54. Scaramuzzino, C. et al. Protein arginine methyltransferase 6 enhances polyglutamine-expanded androgen receptor function and toxicity in spinal and bulbar muscular atrophy. Neuron 85, 88–100 (2015).

55. Arroyo, D. S., Soria, J. A., Gaviglio, E. A., Rodriguez-Galan, M. C. & Iribarren, P. Toll-like receptors are key players in neurodegeneration. Int. Immunopharmacol. 11, 1415–1421 (2011).

56. Marcora, E. & Kennedy, M. B. The huntington’s disease mutation impairs huntingtin’s role in the transport of NF-?B from the synapse to the nucleus. Hum. Mol. Genet. 19, 4373–4384 (2010).

57. Trager, U. et al. HTT-lowering reverses huntington’s disease immune dysfunction caused by NFkB pathway dysregulation. Brain 137, 819–833 (2014).

58. Flood, P. M. et al. Transcriptional factor NF-ĸB as a target for therapy in parkinson’s disease. Park. Dis. 2011, 216298 (2011).

59. Ghosh, A. et al. Selective inhibition of NF-kappaB activation prevents dopaminergic neuronal loss in a mouse model of parkinson’s disease. Proc. Natl. Acad. Sci. U. S. A. 104, 18754–18759 (2007).

60. Pizzorusso, T., Ratto, G. M., Putignano, E. & Maffei, L. Brain-derived neurotrophic factor causes cAMP response element-binding protein phosphorylation in absence of calcium increases in slices and cultured neurons from rat visual cortex. J. Neurosci. 20, 2809–2816 (2000).

61. Zuccato, C. et al. RESEARCH ARTICLE: Systematic assessment of BDNF and its receptor levels in human cortices affected by huntington’s disease. Brain Pathol. 18, 225–238 (2008).

62. Obrietan, K. & Hoyt, K. R. CRE-mediated transcription is increased in huntington’s disease transgenic mice. J. Neurosci. 24, 791–796 (2004).

63. Andersson, M., Konradi, C. & Cenci, M. A. cAMP response element-binding protein is required for dopamine-dependent gene expression in the intact but not the dopamine-denervated striatum. J. Neurosci. 21, 9930–9943 (2001).

64. Devi, L. & Ohno, M. PERK mediates eIF2a phosphorylation responsible for BACE1 elevation, CREB dysfunction and neurodegeneration in a mouse model of alzheimer’s disease. Neurobiol. Aging 35, 2272–2281 (2014).

65. Mantamadiotis, T. et al. Disruption of CREB function in brain leads to neurodegeneration. Nat. Genet. 31, 47–54 (2002).

66. Block, R. C., Dorsey, E. R., Beck, C. A., Brenna, J. T. & Shoulson, I. Altered cholesterol and fatty acid metabolism in huntington disease. J. Clin. Lipidol. 4, 17–23 (2010).

67. Aviles-Olmos, I., Limousin, P., Lees, A. & Foltynie, T. Parkinson’s disease, insulin resistance and novel agents of neuroprotection. Brain 136, 374–384 (2013).

68. Russo, C. V. et al. Insulin sensitivity and early-phase insulin secretion in normoglycemic huntington’s disease patients. J. Huntingtons Dis. 2, 501–507 (2013).

69. Sanders, L. H. et al. Mitochondrial DNA damage: molecular marker of vulnerable nigral neurons in parkinson’s disease. Neurobiol. Dis. 70, 214–223 (2014).

70. Muller, S. K. et al. Lewy body pathology is associated with mitochondrial DNA damage in parkinson’s disease. Neurobiol. Aging 34, 2231–2233 (2013).

71. Moreira, P. I., Carvalho, C., Zhu, X., Smith, M. A. & Perry, G. Mitochondrial dysfunction is a trigger of alzheimer’s disease pathophysiology. Biochim. Biophys. Acta 1802, 2–10 (2010).

72. Coppede, F. An overview of DNA repair in amyotrophic lateral sclerosis. Sci. 11, 1679–1691 (2011).

73. Park, S. G., Schimmel, P. & Kim, S. Aminoacyl tRNA synthetases and their connections to disease. Proc. Natl. Acad. Sci. U. S. A. 105, 11043–11049 (2008).

74. Cha, J.-H. J. Transcriptional dysregulation in huntington’s disease. Trends Neurosci. 23, 387–392 (2000).

75. Hoss, A. G. et al. MicroRNAs located in the hox gene clusters are implicated in huntington’s disease pathogenesis. PLoS Genet. 10, e1004188 (2014).

76. Hoss, A. G. et al. mir-10b-5p expression in huntington’s disease brain relates to age of onset and the extent of striatal involvement. BMC Med. Genomics 8, 10 (2015).

77. Lesage, S. & Brice, A. Role of mendelian genes in “sporadic” parkinson’s disease. Park. Relat. Disord. 18, Supplement 1, S66–S70 (2012).

